# How bibliometric analysis can help to specify and track interdisciplinary research areas: A term-based analysis approach

**DOI:** 10.1101/2025.05.12.653370

**Authors:** Tim Streeb, Christoph Schreyvogel, Christian Jehle, Stefan A. Rensing

## Abstract

Inter- and transdisciplinary research is carried out with increasing frequency and intensity. Bibliometric analysis of disciplinary research is aided by i) classical topical categories that are used by databases cataloguing publications, and ii) via journals that either represent a topic, or attach appropriate disciplinary keywords to papers. In contrast, bibliometric analysis of interdisciplinary research is difficult to perform using disciplinary keywords. Here, we used unigram and bigram term analyses in order to analyse interdisciplinary research areas. These analyses are based on several training datasets representing different research areas at the same institution or with dispersed affiliations. Our results show that using modest investment in query determination, interdisciplinary research areas can be defined and tracked via term-based analysis of their publication output.

## 1. Introduction

Interdisciplinarity and international visibility are becoming increasingly important for larger research associations, especially universities. Bibliometric analyses allow institutions to track the temporal development of research trends, and enable and aid rankings that are tools to measure such visibility. Research conducted within disciplinary frameworks provide the giants’ shoulders on which inter- and transdisciplinary approaches become possible. Incorporating collected knowledge from other fields into one’s own research and thus gaining crossdisciplinary insights is becoming increasingly important in science (Glänzel & Debackere, 2022). Hence, it is mandatory to develop appropriate metrics that allow to perform bibliometric analyses of inter- and transdisciplinary research areas (Leydesdorff & Ivanova, 2021).

Interdisciplinarity arises precisely from overlapping diverse social and scientific fields, exceeding the framework and topic definitions of a single subject area. Characteristics of interdisciplinary research range from the application of the same methods to joint problem solving to close cooperation across several disciplines. The National Academies of Sciences, Engineering and Medicine in the US describes this as follows: “Interdisciplinarity is a mode of research by teams or individuals that integrates information, data, techniques, tools, perspectives, concepts, and/or theories from two or more disciplines […] to solve problems whose solutions are beyond the scope of a single discipline or area of research practice (COSEPUP, 2005).” According to this definition, interdisciplinary research is characterised by knowledge transfer and knowledge integration. Both can take place at different levels (Glänzel & Debackere, 2022).

Disciplinary research is well described by keywords such as hydrology, computational linguistics, or plant genetics, to name a few. However, interdisciplinary research areas are often novel and lack such terminology, or – if they are already established – might be difficult to describe by intersections or unions of disciplinary keywords. This means that bibliometric analysis of interdisciplinary research using classical topic categories of disciplinary research (such as “web of science categories” in Clarivate’s database Web of Science) cannot be used in a straight forward way, since the scope of interdisciplinary research is often just a small part of each contributing discipline regarding all its possible topics and scientific questions, as well as its assigned publications. The classification scheme for subject categories on the journal level (like Web of Science categories) usually cannot be used directly for the interdisciplinary assignment of publications as well as for retrieving them from databases (Xiong & Zhou, 2023). On the other hand, using a classification scheme – the so called Leiden methodology (Waltman & Van-Eck, 2012) – based on direct citation relations between publications requires considerable time investment, at the same time lacking topic-related classification. Bibliometric analysis is not yet widely used for the specification and tracking of interdisciplinary research areas (Donthu et al., 2021). With the help of statistical methods, bibliometrics attempts to offer a quantitative insight into the scholarly/scientific literature and at the same time provide information about the knowledge within a subject area by examining and analysing information recorded in databases, such as citations, references and keywords (Van Raan, 2005). Depending on the information needed for the analyses, differentiated tools such as citation analysis can be used. Keywords in particular, i.e., terms that verbalise the core of a research article, can provide useful information about research topics and trends in a particular discipline and reveal interdisciplinary connections (Leung et al., 2017). Keyword-based approaches for tracking interdisciplinary research areas have been published for areas like synthetic biology, artificial intelligence or nanotechnologies (Liu et al., 2021; Shapira et al., 2017; Suominen et al., 2016). The analysis of most frequent terms in a set of publications of an interdisciplinary research area enables a more specific description of both the content as well as its topical focus than the common keywords of disciplinary research would enable. To distinguish common disciplinary keyword approaches from the approach described here, we are using “term” instead of “keyword” for interdisciplinary bibliometric analyses based on word frequencies.

In this context, this paper aims to (1) give an overview of the bibliometric term analysis using a set of publications of a defined interdisciplinary research area as basis, (2) test how interdisciplinary research areas can be captured using this analysis method and, (3) introduce statistical metrics for defining and evaluating the accuracy of interdisciplinary research areas. As a showcase, we focus on two examples: i) research areas that describe interdisciplinary research at a University and ii) a collaborative funding scheme that by design is interdisciplinary and has dispersed affiliations of its contributing authors.

The methodology presented in this paper focuses on research institutions, collaborative research projects and related stakeholders (e.g. funding organisations) to track the development of an interdisciplinary research area and benchmark against competitors. A better ability to evaluate or qualify interdisciplinary research can aid these stakeholders. This paper is thus positioned in an area of increased interest as institutions recognize the limitations of disciplinary boundaries as well as the difficulties in overcoming these limitations. Thus, the aim of this paper is to perform bibliometric analyses not by using disciplinary keywords but by use of multiple (topical) terms describing the topic of research as defined by a set of representative publications of the interdisciplinary research area. Furthermore, we aim to provide a set of practical guidelines for term selection in interdisciplinary research papers and how to calculate the appropriateness of keywords selected.

## 2. Methodology

### Data sources

The authors acknowledge that open science and FAIR data use are valid goals. Hence, open and transparent databases/aggregators such as OpenAlex (https://openalex.org/) or OpenAIRE (https://www.openaire.eu/) should be supported. For the time being, out of considerations of practicality, the data of the bibliometric analyses performed here are derived from the Web of Science (WoS) Core Collection provided 2023 by © Clarivate (Clarivate Analytics (Deutschland) GmbH, Munich) via access from the German Higher Education and Research Portal (DFN-AAI). Web of Science is one of the largest scientific multidisciplinary databases and contains more than 66.9 million articles from the natural sciences, social sciences, and humanities (Friese & Nuyts, 2017). The broad scope of the database corresponds to the interdisciplinary approach of the analysis, while tools such as PubMed (https://pubmed.ncbi.nlm.nih.gov/) have a more narrow focus (in that case on biomedical literature).

We used four exemplary interdisciplinary research areas for our study. Three are research areas at the University of Freiburg of the period 2018-2023, namely (A) “Epigenetics, Immunology and Cancer Research” which focuses on immunodeficiencies and cancer diseases to gain an understanding of molecular physiology through the elucidation of gene functions towards new treatments, and is complemented and partially overlapping with research into epigenetic control of cell development and cell identity, (B) “Data Analysis and Artificial Intelligence” which is characterised by the integration of machine learning (in particular deep learning) for robotics, computer vision, symbolic AI and data analysis and thus covers all topics of future intelligent systems from automated pattern discovery in data to robotic control and (C) “Complexity of Nature and Future Ecosystems” which focuses on research related to resilience and adaption potentials, as well as the role of biodiversity for the functioning of ecosystems. The fourth example (D) is the DFG-funded (Deutsche Forschungsgemeinschaft) priority programme 2237 “MAdLand – Molecular Adaption to Land: plant evolution to change” which has its focus on the identification of genetic mechanisms underlying adaptive evolution of plant morphology, physiology, biochemistry, cell biology and biotic interactions in conjunction with plant terrestrialization.

The analysis timeframe was restricted to the period between the formation of the research area (2019)/priority programme (2020) and the time at which the initial analysis was conducted (2022). For the specification of the research areas, principle investigators of the respective area were asked to name a maximum of three of their publications which they consider most relevant for the interdisciplinary area. This resulted in 60 publications in area (A), 42 publications in area (B), 47 publications in area (C) and 40 publications in area (D). In addition to the association with the respective area and the author’s name abbreviation in WoS, the title and abstract of the publications were recorded for each of these (Table SI1). The latter forms the data basis for defining the research areas with the help of terms present in the relevant publications using Web of Science as a repository.

### Word frequency analysis to determine terms

In order to perform word frequency analyses, the 24,495 words that were used in the abstracts of the 189 publications were assigned to their respective research area by importing the abstracts into a spreadsheet (Microsoft Excel 2019) and annotation according to research area affiliation. For the subsequent word count analysis, the abstracts of each research area were transferred to the WordClouds tool (https://www.wordclouds.com/) resulting in a data set listing the frequency of each word for each research area (WordClouds.com, 2022) If words differ in their singular/plural form or share the same word stem but differ in their ending, they were recorded and counted as separate words. For the analysis of bigrams, the Bibliometrix R-Tool version 4.0 and its Shiny Platform was used (www.bibliometrix.org; (Aria & Cuccurullo, 2017), R-version 4.2.2 (2022-10-31)).

Subsequently, the words were manually screened to determine whether they might be considered typical for the description of the research areas and hence represent appropriate terms to describe the interdisciplinary research area. Words, in particular verbs such as “can”, “show”, “use” etc., which are essential for the formulation of grammatically correct sentences, but which are not directly related to the research area or are not used exclusively in this field, were not considered.

Based on the initial sorting, the absolute frequencies of the terms were determined in the next step. A threshold value was then defined, which determines the minimum number of times a word has to be mentioned in the abstracts to be considered characteristic of the research area. This threshold was defined as the 95% quartile, which comprises the 5% most frequent words.

However, to be considered characteristic of the research area, terms must not only occur frequently overall (frequent words), but they also need to be included in many different publications/abstracts (broadly used words). An example of this in the research area (C), Complexity of Nature and Future Ecosystems, is beech, which occurs frequently in the abstracts with a total of 19 mentions, but at the same time is only mentioned in 5 different abstracts, i.e. in 11% of all abstracts (Fig. 1b). Beech was therefore not used as a term for the query.

Hence, the relative (%) frequencies were used to avoid the overweighting of terms that, despite their high absolute frequency, appear in only a small number of abstracts. Terms that are both frequently used in absolute and relative numbers are considered suitable for describing the research area.

A high threshold level, such as 30% for the relative frequency of terms, leads to the inclusion of only those terms that are particularly prevalent in the characterization of the research area. This typically results in a low false discovery rate (FDR) of a query consisting of these few but specific terms, but also low sensitivity (cf. Results). Alternatively, a lower threshold level, such as 15%, results in a broader definition by including a larger number of terms in the characterization of the research area (Fig. 1b); such a query is generally more sensitive but has a higher FDR. The minimum, maximum and average values for absolute and relative frequencies for research areas A-D, together with the values for each term, are shown in Fig. SI1.

Ultimately, defining the threshold levels depends on the FDR/sensitivity balance that shall be achieved.

Following this procedure only a few potential terms remained that should best verbalize the core of the research area (see Figure 1a and 1b).

**Figure 1a:**
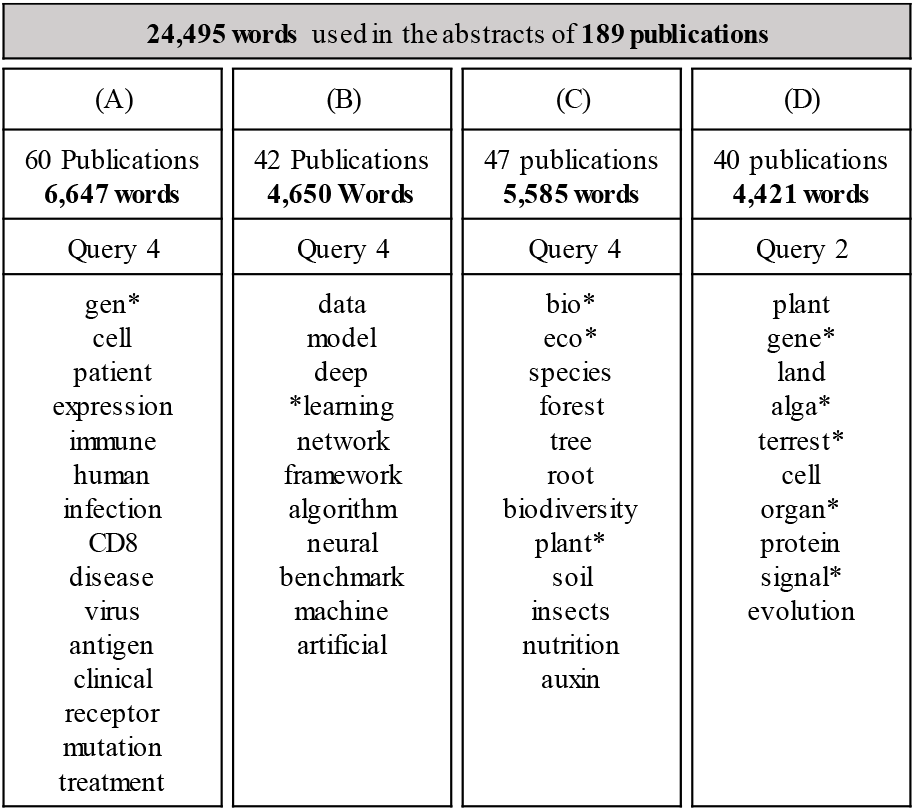
Terms used for the characterization of each research area for the best performing query.

**Figure 1b:**
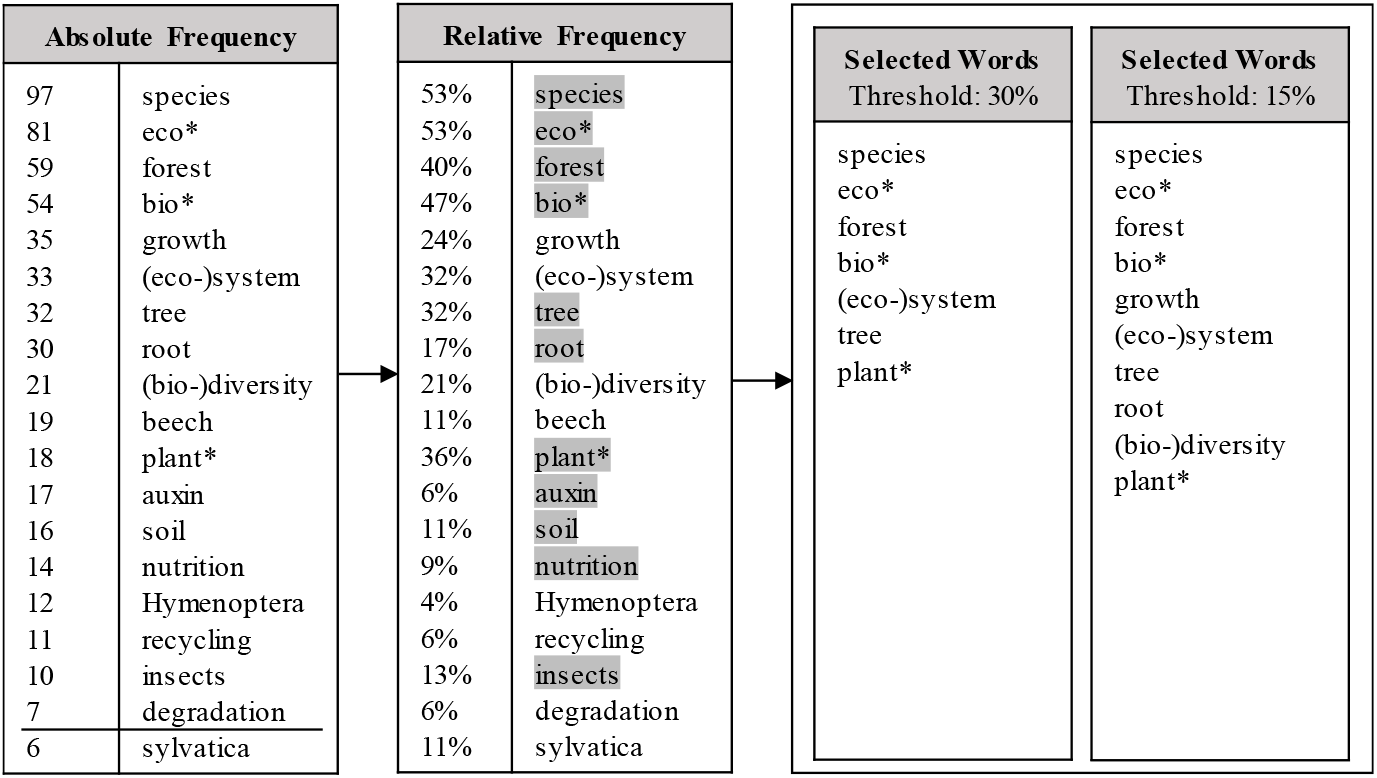
Exemplary illustration of the selection of terms that are used to define research area (C) “Complexity of Nature and Future Ecosystems”. The 95% quartile (7) is shown as a horizontal line in the absolute frequency box; absolute frequency is derived from the training data set via word count analysis. Relative frequency describes in which % of training set abstracts the term occurs. Terms used in Area (C) QUERY 4 (Table 4) are highlighted in grey in the relative frequency box; note that (eco-)system and (bio-)diversity are covered by eco* and bio* and are used as mandatory AND terms, while all the other terms are used with OR flags.

### Query formation

Subsequent bibliometric analysis was performed using the “Advanced Search Query Builder” in Web of Science, allowing to filter and analyze articles based on a wide range of information defined by field tags such as TI (titles), AB (abstracts), AU (authors), PY (year of publication), TS (topic terms), where the latter combines a set of fields (including TI, AB as well as Author Keywords and Clarivate’s Keywords Plus®). Multiple field tags can be linked with search operators to combine terms to broaden or narrow the search (Clarivate Analytics, 2020). The Boolean operator AND can be used to include two or more search terms that must be found. For example, it may be useful to combine related terms such as *tree* AND *forest* to exclude publications from unwanted areas, such as statistics (e.g. tree-shaped diagrams). In contrast, OR can be used to find results with one or the other search term, which in turn helps to connect different subject areas and to pick up on the interdisciplinary approach. The wildcard character * (representing 0..n characters) offers an additional way to refine the search query by allowing different spellings of the same term in the search and/or allowing WoS to find terms with the same stem (e.g., bio* will find publications containing the term biology as well as bio-diversity etc.) (University of Hawai’i at Mānoa, 2022).

Field tags, Boolean operators, and the terms potentially describing the research area were thus combined to form queries, providing the basis for further analysis. In the case of “Complexity of Nature and Future Ecosystems” the query looks like the following, for example:

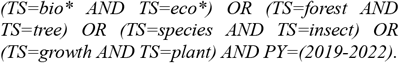

The four brackets represent the different characteristics (related to the disciplinary parts) within the research area, each of them connecting two terms using AND, while being separated from each other with OR. The first bracket is defined in broader terms by using the wildcard character *. Linking the term stems *bio\** and *eco\** seemed to make sense because both terms are thematically very close and jointly seem to specify the research area very well. On the other hand, it does not seem to make sense to link the term(-stem) *eco\** and *growth*, for example. Due to the proximity of *eco*\* to terms of the area of *eco*nomics, especially in connection with *growth*, the results of the query would be misleading. Following this logic, the remaining brackets were created accordingly until all the relevant term were accommodated in one query. The same was repeated for the other research areas. Know-how of the research area in question is helpful to improve queries. In our approach, initial results were presented to each of the four research areas, and feedback provided to further improve the queries.

### Classification

The confusion matrix used for classification problems is a table with two rows (representing the actual condition) and two columns (the prediction). It reports the numbers of true positives, false positives, true negatives and false negatives. From such a matrix, many useful statistics such as sensitivity (the ability to find many true positives) or specificity (the ability to exclude false positives) can be computed.

Within this context, the population (U), positives (P), negatives (N) as well as true positives (TP), false negatives (FN), false positives (FP) and true negatives (TN) must be computed or estimated. For example, for a University research area we consider the population (U) as the total number of articles retrieved with the PI’s names of the research area in the selected time period (2019-2022), but without filtering for the correct institutional affiliation. All publications of the research areas’ PIs that can be associated with the institution (here: University of Freiburg (UFR)) via a given WoS affiliation query are counted as a positive (P), while publications that cannot be attributed to the University of Freiburg are referred to as negatives (N), thus reflecting the general need for consequent usage of unique author identifiers. This definition is possible if none of the PIs moved to or from the University during the time period analysed, which was the case for A, B and C here. A publication is considered TP if it can be associated with the UFR as well as with the PIs of the research area (= actual condition), and at the same time is retrievable with the query including the chosen terms (= prediction). A publication is considered FN if it can be associated with the UFR and with the PIs of the research area, but is not retrievable with the query. Accordingly, a publication that can be found with the query but cannot be assigned to a PI of the research area is considered FP. We see no way to sensibly define N or TN.

In the case of research area D, the priority programme MAdLand, FP could not be determined. In the case of the University research areas, we are narrowing down the search results in WoS using the query in such a way that only those publications are listed that have an affiliation with the preselected organisation (here: UFR) in the analysed time period. In the next step, the publications that can be assigned to the PIs of the research area are subtracted from those results. The remainder are all publications by persons who publish with the affiliation UFR, who are found with the query, but who are not part of the research area (= false positives). No such unique affiliation exists in the priority program MAdLand, and the relatively high number of institutions with which the PIs are affiliated would make a corresponding analysis awkward. Hence, in case of MAdLand the statistical indicators are defined as follows: TP as number of publications by MAdLand PIs found with the query; FN as number of publications by such authors not found with query; FP as publications found with the query that cannot be assigned to PIs of the priority program.

Based on these indicators, we can calculate the sensitivity or true positive rate (=TP/(TP+FN)) and the false discovery rate (FDR) (=FP/(FP+TP)), which can be used to assess how well the respective research area is specified based upon the chosen terms.

## 3. Results

### Classification based on the confusion matrix

In order to determine the quality/accuracy of the term-based query approach, we decided to use the classical confusion or error matrix approach commonly used for classification problems (Stehman, 1997). While the principal considerations and setup are described in Methodology, we will point out here potential caveats in the definition of categories.

For selecting good queries, a result with high sensitivity and low FDR would be favorable; specificity (=TN/(TN+FP)) was not be calculated since TN could not be defined (see Methodology). FDR was used as an alternative to judge how specific the query results are. For example, for a University research area a result with 100% sensitivity and 0% FDR could be interpreted as follows: All publications of the authors were found with the query and are affiliated with the UFR, no further publications can be found with the given query that cannot be assigned to the research area. In reality, this is almost impossible, which is due to the fact that there are almost no words, or terms, that are exclusive to one disciplinary or even interdisciplinary scientific field. In other words, due to the nature of inter- and transdisciplinary research it is impossible to separate different research areas strictly from each other as there exists a lot of overlaps regarding topics, scientific questions, methodology and so forth. Also, authors will publish in their own disciplinary as well as in other interdisciplinary research areas. Hence, achieving both a high sensitivity and a low FDR is very difficult.

### University research areas (known single affiliation of PIs; A, B, C)

We will first take a closer look at a university research area. The considerations and results obtained can be easily transferred to other such areas.

The analysis of the research area (C) “Complexity of nature and future ecosystems” is based on 47 publications by 16 PIs (Table SI1). The period of analysis is limited to the years 2019-2022. In this period, a total of 1,427 publications (U) can be found, of which 385 publications (P) can be assigned to the University of Freiburg (UFR) and 1,042 publications (N) have no relation to the UFR (these are primarily based on redundant author name matching). Based on the respective query including the chosen terms for this research area, the indicators TP, FN and FP are derived as outlined above (Figure 2).

**Figure 2:**
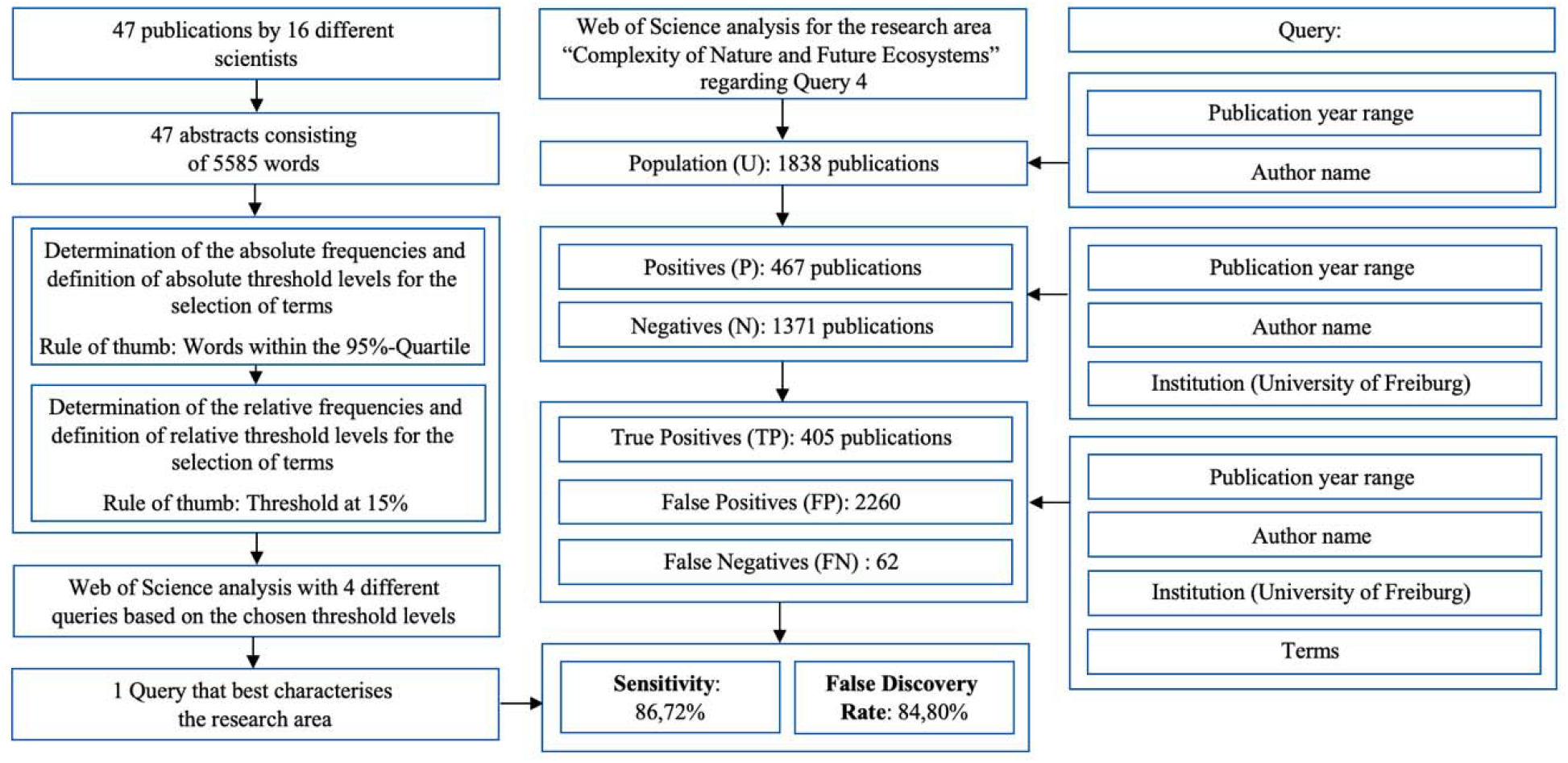
Flow chart of bibliometric analysis including of the example research area (C) “Complexity of Nature and Future Ecosystems”, QUERY 4.

QUERY 1 is defined as follows:

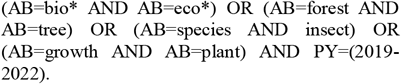

This query consists of four topic-specific brackets, which aim to represent the diversity of the research area, and the tailing topic-independent bracket, which is used to restrict to the period of publication. The first bracket serves to broadly describe the research area. The terms bio* and eco* were chosen because they are both very common among the abstracts of the area. The second bracket uses the terms forest and tree to describe the topic of forest science, which plays a major role within the research area. The third bracket takes up this aspect again in a different way and focuses on the animals studied. The fourth topic-specific bracket completes the description of the research area and serves to cover publications that have not yet been addressed by the previous topic fields. The common feature of the brackets is that they use the “field tag” AB (abstracts), i.e., only publications that contain the terms in their abstracts are found. The idea behind this is that an abstract analysis, compared to a full-text analysis (see QUERY 3 and QUERY 4 later in the text), should provide a lower FDR, since terms used in the abstracts might be particularly relevant for the description of the publication and thus the research area.

The following values were determined for QUERY 1. 139 publications were found and can be assigned to the authors of the research area, and are also connected to the UFR (TP). Consequently, 246 publications (P 385 – TP 139; FN) by authors with UFR affiliation could not be found using the query. 301 publications (FP), on the other hand, were found with the query, but could not be assigned to the PIs associated with the research area. By implication, these 301 publications might originate from authors who are not (yet) known as contributors to the University research area, but who actually do publish in this same area. Alternatively, and more likely, many of these publications might have no linkage at all to this interdisciplinary research area, because the query was not specific enough. The sensitivity of QUERY 1 was 36.41% and the FDR 68.41%.

QUERY 2 is defined as follows:

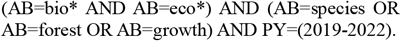

It differs from QUERY 1 insofar as it contains only two topic-specific brackets linked with AND, and the terms within the second topic-specific bracket linked with OR. This leads to a required relationship between the first bracket, which roughly describes the research area, and the second bracket, which lists the most relevant terms of each topic within the research area. Compared to QUERY 1, which already returns records if one of the subject areas is listed in the publication, QUERY 2 only returns records if the conditions from the first bracket are met and at least one other term from an additional subject area has been used. Intuitively, the QUERY 2 defines the research area more strictly, which should lead to a decrease in the FDR but also the sensitivity. The trade-off between the two ratios is illustrated using the calculation “(Sensitivity-FDR)+1” (Figure 3). Queries with a higher value for TP and a lower FDR show a better balance and are preferable. Since the maximum score that can be achieved is 2 (200%), values below 1 (100%) need to be regarded with caution (cf. Figure SI2). It should be noted that the underlying population for TP and FDR differs and hence this calculation is inferior to a sensitivity/specificity balance. For the above-mentioned QUERY 2, a sensitivity of 19.74% and a FDR of 59.14% was determined, i.e., our assumptions that both FDR and sensitivity are lower compared to QUERY 1 are confirmed. Due to the very low sensitivity, the balanced performance of QUERY 2 is worse than QUERY 1 (grey dots in Figure 3).

**Figure 3:**
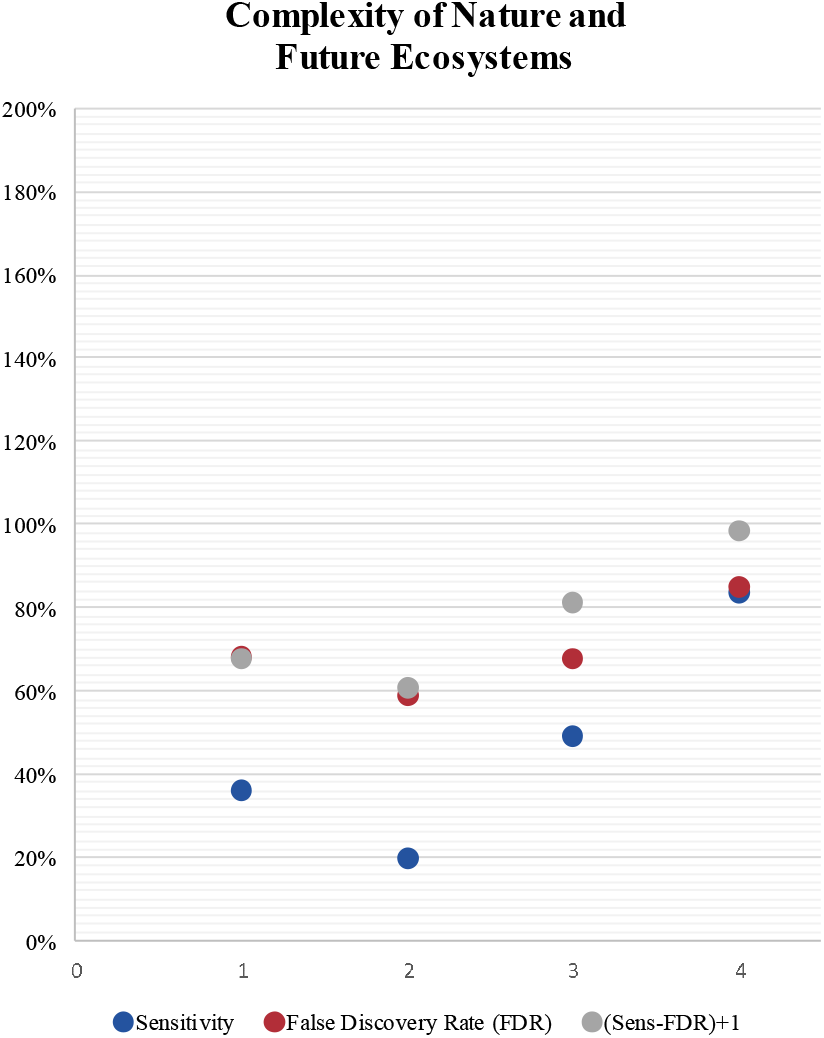
Sensitivity, False Discovery Rate (FDR) and (Sens-FDR)+1 of the research area (C) “Complexity of Nature and Future Ecosystems” for each QUERY 1-4.

QUERY 3 is defined as follows:

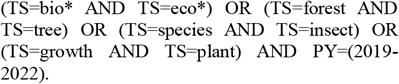

This query does not differ in its structure from the QUERY 1. The only difference is that by using the field tag TS (“topic terms”) the query takes place at topic level, i.e. for the search WoS adds the title, the author keywords and the “KeyWords Plus®”. KeyWords Plus are words or phrases that frequently appear in the titles of an article’s references, but do not (necessarily) appear in the title of the article itself. Using this QUERY 3, sensitivity is 49.09% and FDR 67.91%. If we compare these values with the QUERY 1, we can see an increase in sensitivity of 13 %, while at the same time the FDR has slightly increased. The balanced score (Figure 3) is superior to QUERY 1 and 2. Using this third query, we find more publications overall and more publications with affiliation Freiburg due to the larger search spectrum of the field tag “TS” compared to the field tag “AB”. This results in an overall increase in sensitivity. At the same time, however, the larger search spectrum leads to more publications being found by the query that cannot be assigned to the authors of the research area (FP). In effect, this query performs better than 1 and 2 (Figures 3, 4).

**Figure 4:**
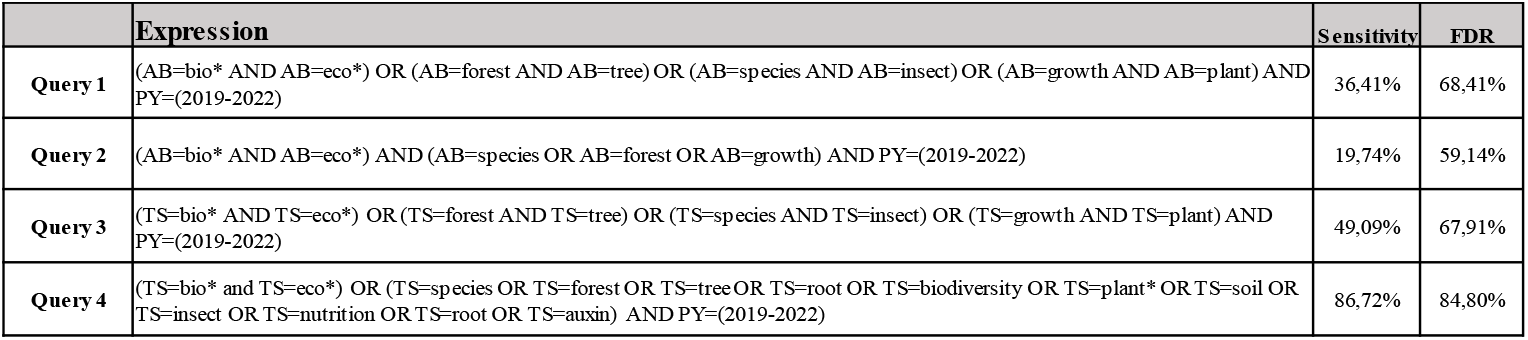
Sensitivity and False Discovery Rate of the research area (C) “Complexity of Nature and Future Ecosystems” queries.

QUERY 4 is defined as follows:

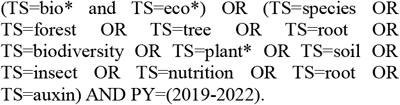

It is thus similar in structure to QUERY 2, with its two topic-specific brackets, but here the two topic-specific brackets are not connected with AND but with OR. Another difference to the second query is that here, analogous to the third query, the field tag “TS” was used. Also, eight terms were added to the second topic-specific bracket, which consists now of thirteen terms. By combining these expanded term list in the second bracket with OR statements, we attempt to group similar terms because not every term appears in every abstract. The terms chosen for QUERY 4 all have an absolute frequency in the top 5% quartile (average 37.42) and a relative frequency of 9% or higher (average 28.17%; Fig. 1b, Fig. SI1). Taken together, these measures lead to a further strong increase of sensitivity to 86.72% at the price of an increase of FDR to 84.80%. While this query shows the best sensitivity of all queries (Figure 3), the high FDR renders it undesirable to perform further downstream analysis like ranking of various institutions. In Figure 4, the values for sensitivity and FDR are summarized for the four queries for the research field (C). Similar diagrams are shown in Figure SI3 and SI4 for the research areas A and B.

### Research area with dispersed PI affiliations (D)

These findings outlined above cannot be transferred 1:1 to the MAdLand setting (D), primarily because no single affiliation can be used to estimate the FDR. However, three different MAdLand datasets allow the calculation of the sensitivity over the time course. The first dataset encompasses the period 2011-2020 (publications that thematically belong to MAdLand and were published by MAdLand PIs, yet before the start of the program; preliminary work). The second set comprises the period 2021-2022 (publications that have been published during the first funding round of the program), while the third set comprises the entire period 2011-2022 (Table SI1+2). One might expect that relevant terms appear with a higher probability in the publications of the program than in previous work, and thus a sharpening of the profile is expected to take place that might be reflected in the sensitivity.

In light of the fact that no FDR can be calculated for MAdLand, the question remains as to how the query can be assessed without this indicator. We obtain an answer by looking at the second and fourth MAdLand queries. These are similar to the previously explained fourth query of research area C, they differ from each other only in that the two topic-specific brackets are linked once with OR (QUERY 2) and once with AND (QUERY 4).

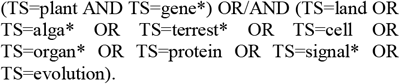

Hence, QUERY 2 will detect a publication if it either includes the two terms from the first topic-specific bracket OR a term from the second topic-specific bracket. In combination with a very broadly defined second bracket or terms that may also be characteristic for other research fields, such as *cell* or *organ*\*, this leads to numerous publications being found that are not relevant (FP; for example from medical research). The situation is different, however, if the two topic-specific brackets are connected with AND: many publications that are not part of the research area can be excluded. In effect, a query in which the topic-specific brackets are connected with OR delivers a high sensitivity, while at the same time collecting many FP. The reverse is true for a query in which the topic-specific brackets are linked with AND. This yields a lower sensitivity while at the same time the profile field is expected to be specified more precisely. As expected, the sensitivity of QUERY 4 is a lot lower than for QUERY 2 for all three different datasets of the MAdLand setting (Figure 5). For the evaluation of the priority program – contrary to the research areas (A), (B) and (C) – due to the dispersed affiliation we use the ORCID-IDs of the principal investigators rather than the affiliation to restrict to the (D) publications. Obviously, this procedure can only be applied if the PIs use their ORCID-IDs in their publication. Although we cannot calculate the FDR, QUERY 4 in turn is expected to be more specific – as evidenced by the lack of medical topics in the WoS Citation Topics Meso (Table SI3). Furthermore, within both queries the sensitivity is highest for the time period after the start of the program, and lowest for the period before the start. Hence, as expected, the actual publications from the field reflect the query better / can be found with higher sensitivity than those of the same authors before the start of the program.

**Figure 5:**
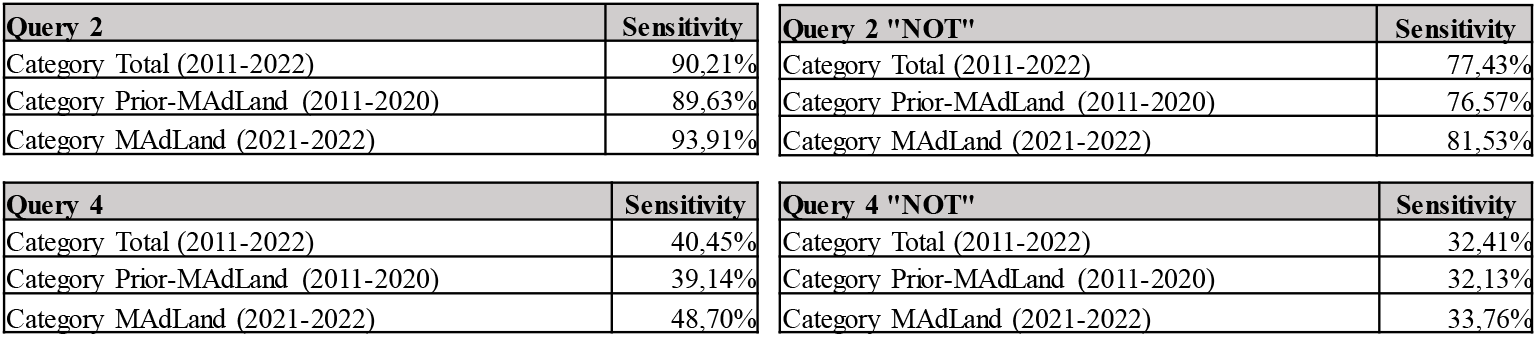
Trends in Sensitivity for the priority program MAdLand regarding QUERY 2 and QUERY 4 with unigrams

### Ranking

An alternative to get a basic idea of the specificity of the queries, even without calculating the FDR, is the analysis tool provided by WoS. This tool processes all publications and lists the results in descending order in various categories. With regard to the analysis of the queries of the research areas, the category “Affiliation” is particularly useful for us, as it assigns the publications retrieved to their affiliation to the respective institution. If this result is exported, further filtering possibilities arise. One example, detection of (or lack of) institutions not expected may be used to estimate the specificity of the query in absence of FDR calculation. In addition, apart from the evaluation of the quality of the respective query, the category “affiliations” can be used to sort e.g. on national level to rank institutions in terms of how much they contribute to an interdisciplinary research area. An example for such a ranking for the research area (C) “Complexity of Nature and Future Ecosystems” on the national level in Germany, performed with the above-mentioned QUERY 4, is shown in Figure 6. Based on the sensitivity and FDR, the bar for University of Freiburg consists of ca. 300 TP and ca. 1,500 FP. It is reassuring that despite the collection of many FP the University of Freiburg is found on the third rank in Germany, an expected result since not all universities will perform in this interdisciplinary research area. Also, the institutions ranked 1^st^ and 2^nd^ as per their webpages pursue related research areas (Göttingen: Sustainable Land Use and Bioversity; TUM: Sustainable Living Environment).

**Figure 6:**
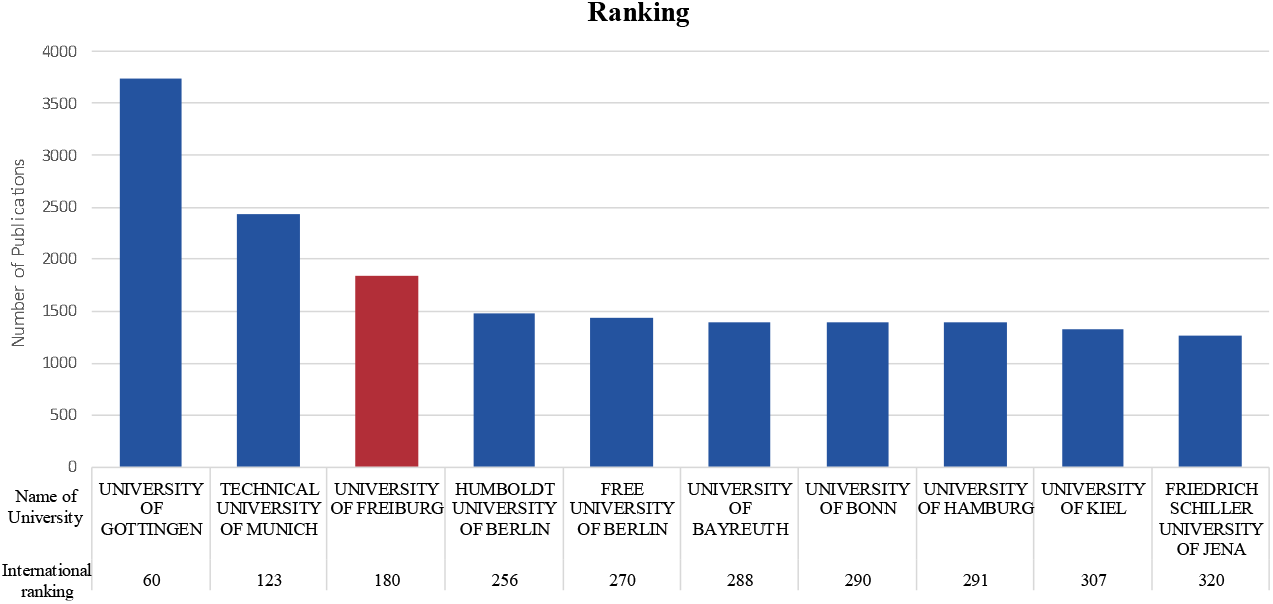
Institutional ranking by affiliation as recorded in WoS, based on QUERY 4 which characterizes best the research area (C) “Complexity of Nature and Future Ecosystems”

### Bigram analyses

In analogy to the analyses performed with unigrams (one-word terms) for the priority program (D) “MAdLand” we also applied bibliometric analysis with bigrams, i.e. with the most relevant terms consisting of two words. For the term count analysis of the bigrams, we first retrieved all publications from the Web of Science repository by entering all DOIs (Digital Object Identifiers) of each MAdLand publication (Table SI1). Then, this set of publications of the priority program was transferred to the Shiny tool of the Bibliometrix platform as a text file containing information about title and abstracts of the publications. A subsequent analysis resulted in a data set listing the frequency of each bigram (WordClouds.com, 2022). If the terms differ in their singular/plural form or share the same word stem but differ in their ending, they were recorded and counted as separate terms. Subsequently, the terms were manually screened to determine whether they might be considered typical for the description of the priority program and hence represent this interdisciplinary research area. The preliminary sample consisted of approximately the 100 most frequent terms. Based on these, a further manual screening was conducted. The selected most relevant bigrams for MAdLand are: *land plant, plant lineage, flowering plant, streptophyte alga*, morphological complexity, specialized metabolite, terrestrial habitat, abiotic stress, common ancestor, gene expression, green lineage, oil body, phenotypic plasticity, plant species*.

It should be noted that for bigrams it is difficult to apply these criteria, as the absolute frequencies are often much smaller. Thus, one has to select terms which seem to be most relevant for the research area to be investigated and construct the subsequent query in a different way as described in the following.

We applied three different queries, two of which have a similar structure to QUERY 2 and QUERY 4 described for the unigrams (Figure 4).

The MAdLand QUERY 2.1 is defined as follows (Figure 7):

**Figure 7:**
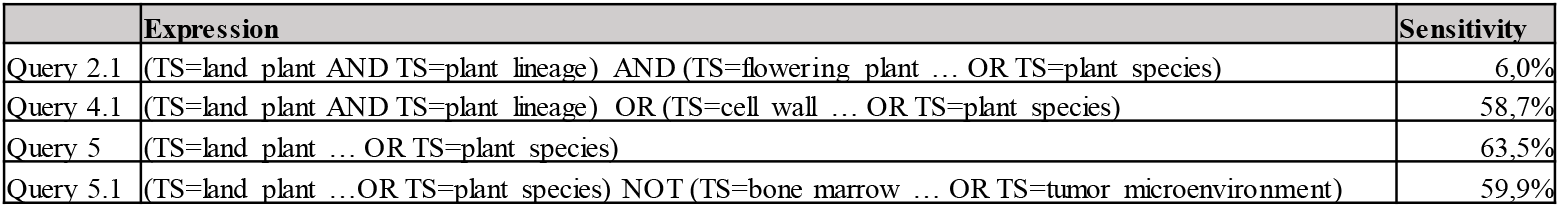
Trends in Sensitivity for the priority program MAdLand regarding QUERY 2.1, 4.1, 5 and 5.1 with bigrams.\

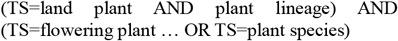

while the MAdLand QUERY 4.1 is defined as follows:

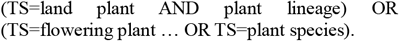

Where the dots in the second bracket indicate all the selected most relevant bigrams for MAdLand which were mentioned before. In case of QUERIES 2.1 and 4.1, the two topic-specific brackets are linked either with AND (QUERY 2.1), respectively OR (QUERY 4.2). The (similar) implication of these queries for the search results in Web of science is mentioned earlier in the text in the context of the unigrams.

For QUERY 2.1, the resulting number of publications retrieved from Web of Science is 15 (TP), far below the number of publications generated in the priority program in the same period (44). The calculated sensitivity is thus very low at 6% (Figure 7, Table SI4). The reason for this is that, contrary to the unigrams, the frequency of the bigrams is both lower overall and often found only in a few publications of the program. As mentioned before in the context of the unigrams, a prerequisite for a successful query is that the selected relevant terms occur in many or ideally all publications. Furthermore, as the search query is even more specific by using the AND operator, we cannot track the research area with this query structure. Therefore, the next logical step is to broaden the query by using the OR operator between the first and second topic-specific bracket (QUERY 4.1). The resulting sensitivity is, as expected, higher with 58.7%, detecting 148 TP.

To increase the sensitivity, the query can be defined even broader by combining all the most relevant bigrams exclusively with the OR operator (QUERY 5).

The MAdLand QUERY 5 is defined as follows:

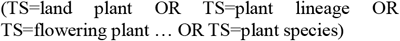

Where the dots in the second bracket indicate again all the selected most relevant bigrams for MAdLand. The resulting sensitivity is slightly higher (63.5%, 160 TP), but it is still lower than for QUERY 2 with the unigrams (cf. Figures 5, 7). Unigrams have a higher sensitivity due to their more general nature. The sensitivity of a search query with bigrams is thus lower because the terms are often more subject-specific and occur with a lower absolute and relative frequency.

As QUERY 5 defines the research area broadly and thus not specifically enough, there are (many) publications that are not part of the program and thus considered as FP. A possibility for excluding these publications is to identify bigrams that do not belong to the research area. Ideally, for the identification of these bigrams, a large set of publications retrieved from a query should be analyzed. As a query for retrieving TP for a defined interdisciplinary research area on a university level often results in relatively low number of publications which are in the order of 100 publications or even lower, we have to construct a query which retrieves more publications. This can be achieved by using a query to retrieve all respective publications on the national or even international level for example (not university level/program level). In the case of QUERY 5, we retrieved 28.121 publications on the national level in Germany for the same period. This large set of publications was transferred (only partially due to size restriction) from Web of Science to the Shiny tool of the Bibliometrix platform as a text file containing information about title and abstracts of the publications. By analyzing the most frequent terms of these publications, we could identify not only the most relevant bigrams (of the research area) as described before, but also thematically wrong bigrams based on manual selection. A manual selection of the resulting bigrams defining the wrong publications comprised terms such as: *bone marrow, human liver, cancer cell, immune system, immune response, inflammatory response and therapeutic target*, to name a few (Table SI4). These bigrams describe medical research and are definitely out of the scope of the program. It must be noted that a manual selection of both the thematically correct and wrong terms requires some knowledge about the research area to be investigated. With the help of these bigrams, defining both the relevant terms of the program as well as the thematically wrong terms, we could optimize the query, which is as follows:

The MAdLand QUERY 5.1 is defined as follows:

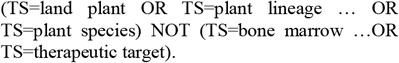

By adding the NOT operator after the topic-specific bracket describing the priority program, we can exclude publications containing the terms listed in the subsequent bracket, which are the abovementioned thematically wrong bigrams. For QUERY 5.1, containing 31 expressions in the subsequent bracket of the NOT operator, the resulting number of publications retrieved from Web of Science is 19,936. This is 41.1% less compared to the number of publications retrieved with QUERY 5 on national level in Germany while at the same time the number of TP is almost the same, i.e. 151 (for QUERY 5.1) and 160 (for QURY 5) on program level. The sensitivity is 59.9% for QUERY 5.1, only slightly lower than for QUERY 5 with 63.5%. As mentioned earlier, it is not possible to calculate FDR directly, but in light of the results for QUERY 5.1, we presumably could reduce the FDR considerably by almost preserving the sensitivity and the number of TP on the program level and at the same time strongly reducing the number of FP, i.e. the thematically wrong publications on national level. Figure 7 summarizes all relevant results of bigram QUERIES 2.1, 4.1, 5 and 5.1. As outlined above, QUERY 5.1 gives the best balanced result. This method – excluding wrong publications with the NOT operator – can also be applied for queries with unigrams. For example, if we add the NOT operator to the original unigram QUERY 2, sensitivity drops from 93.91% to 81.53% (Figure 5) - an acceptable reduction given the fact that medical literature is avoided (in comparison, the strict AND operators in QUERY 4 led to a sensitivity drop to 48.70%).

In summary, a query combining the most relevant bigrams of the program with the OR operator as well as adding a NOT operator to exclude thematically wrong bigrams is a means to both increase the sensitivity and to exclude FP. This method is an alternative to finding the right combination of the most relevant terms with the AND as well OR operators like in MAdLand QUERIES 1 - 4 for the unigrams as shown in Figure 4 as well as QUERIES 2.1, 4.1 and 5 for the bigrams as shown in Figure 7. This method might be applied in cases in which both the absolute as well as the relative frequencies of the terms in the publications are low. Furthermore, the application of this method is suitable for optimizing a query in an iterative way, i.e. after each query the remaining thematically wrong terms could be identified additionally and added iteratively to the subsequent bracket of the NOT operator until the number of retrieved results of a query converges to a stable output or to an acceptable minimum of wrong publications. After this iterative optimization of a query, the quality of the publication output can be assumed to be sufficient for performing evaluation of the research area on e.g. national or international level. The affiliations can then be sorted on national or international level to rank institutions in terms of how much they contribute to an interdisciplinary research area.

### Random sample evaluation

For research areas (C) and (D) we generated random output samples by selecting the 50 most recent publications detected by a given query. Using expert know-how on the respective research area, the TP were determined in these lists of publications.

In area (C), the unigram queries, conducted without restricting to the list of PIs, led to 7 TP among the 50 most recent papers delivered by the best performing QUERY 4 (Figure 3). As for QUERY 4 the sensitivity is 86.72% and FDR 84.80%, we should expect 42.5 FP among a 50 paper sample. Hence, the 7 TP found mirror expectation.

In area (D), for the unigram queries, we identified 3 TP for the best performing QUERY 4 (Figure 5).

For QUERY 4 the sensitivity is 48.7%. Given the 3 TP mentioned above, the 47 FP correspond to an FDR of 94%. Adding the NOT terms to avoid medical research led to 3 TP as well, not (overly) reducing sensitivity, as expected (Figure 5). Additional terms based on expert review of the random sample were added to avoid terms related to non-biological research topics (Table SI4). These lead to an additional reduction of hits (and thus lower FDR/higher specificity), yet at a slight loss of sensitivity.

For the MAdLand (D) bigram analyses, restricting the output to the list of PIs via their ORCID, 15 TP each were detected using QUERIES 4.1, 5 and 5.1. For those queries, the expected number of TP based on their sensitivity in the raw/training data – presented before in the section bigram analyses - is 29 (QUERY 4.1), 32 (QUERY 5) and 30 (QUERY 5.1). Hence, the 15 TP each in the random samples show a twofold under-prediction in real data. This is not a bad result, since a significant portion of TP is indeed detected. Yet, one needs to be aware of the fact that the results on real data are not as good as on training data.

QUERY 2.1 detected 14 publications, all of them determined to be TP. This result reflects a very high specificity of query 2.1 (Figure 7).

These evaluations of the queries show that in general they perform as expected, i.e. that considerations based on the raw/training data may be extrapolated.

### Bibliometric analysis of interdisciplinary research using classical topic categories of disciplinary research

Above we presented a methodology to specify interdisciplinary research areas using terms which appear in publications of areas (A), (B), (C) and (D). As already mentioned in the introduction, bibliometric analysis of interdisciplinary research using classical topic categories of disciplinary research cannot be used in a straight forward way, since interdisciplinary research combining these disciplines is often just a small part of each contributing discipline regarding all its possible topics and scientific questions, as well as its assigned publications. In the following we demonstrate as a showcase for the research area (D), the priority program MAdLand, that these classical topic categories are indeed not suitable to define the exact content of this research.

An analysis of the publications of the research area (D) by entering its DOIs (Digital Object Identifier) in Web of Science led to the assigned topics in the following categories, where the number in bracket is the number of publications assigned:

- Web of Science Categories:

Plant Sciences (26), Cell Biology (5), Biochemistry & Molecular Biology (4), Biology (3), Reproductive Biology (2), Developmental Biology (1), Ecology (1), Evolutionary Biology (1), Genetics & Heredity (1), Multidisciplinary Sciences (1)

- Citation Topics Meso:

Crop Science (22), Photoproductivity (3), Phylogenetics & Genomics (2), Lysosomal Storage Disorders (1), Bacteriology (1), Molecular & Cell Biology – Genetics (1), Animal Sensing (1), Soil Science (1)

- Citation Topics Micro:

Arabidopsis (15), Pectin (4), Jasmonic Acid (3), Photosystem (2), Ceramide (1), Outer Membrane, RNA-Seq (1), Microalgae (1), Hydra (1), Phosphorous (1), Ferns (1), Mitochondrial Genome (1)

According to the definition, Web of Science Categories are assigned at the journal level, i.e. all items in a journal will be assigned the Web of Science Categories of the journal it is published in. Each journal can be assigned a maximum of 6 Web of Science Categories. Contrary to that, the publications are assigned to only one Citation Topic Meso/Micro. Citation Topics represent groups of papers related to one another via citation, i.e. represent a cluster of publications with a defined topic. The level Meso/Micro is a result of the granularity of the clustering scheme.

By looking at the Web of Science Categories assigned to the publications of the priority program, many categories mentioned (such as Plant Sciences, Cell Biology, Biochemistry & Molecular Biology, Developmental and Evolutionary Biology) are plausible and reflect disciplines involved in this interdisciplinary research. Nevertheless, these categories cannot be used to perform bibliometric analysis of the defined interdisciplinary research topic of this priority program, as the topical scope is just a small and specialised part of each contributing discipline. For example, Plant Sciences encompass all analyses of plants, while the focus of the program is evolutionary analyses of particular plant lineages. Hence Plant Sciences allows for a good sensitivity but bad specificity.

This becomes worse for the Citation Topics Meso assigned to the publications of the priority program, since in the strict Sense no single paper is actually assignable to the most frequently found term Crop Science.

Similarly, Citation Topics Micro lists Arabidopsis as the most frequent term, although by definition none of the papers conducts work on Arabidopsis as such, but uses it for comparative purposes.

In summary, these classical topics representing the different disciplines can be used to analyse e.g. its development within an institution and benchmark with competitors. They cannot be applied for specifying interdisciplinary research areas without developing methods to weight individual contributions of disciplinary topics.

### Query generation using a large language model

An alternative approach to generating search queries was investigated using ChatGPT 4.o (accessed 18.10.2024). The model was asked to name the ten most important publications in artificial intelligence in 2023. It was then asked to determine the most frequent words that appear in these ten publications (and to disregard words such as prepositions that appear in any text). As a third step, the model was asked (based on the previous answer) to suggest a list of max. 10 terms that appear specifically in publications about AI (and that the terms should appear in many or most of the ten publications named before) (cf. Table SI5a). The resulting list of top 10 terms (1. Model, 2. Learning, 3. Ethics, 4. Alignment, 5. AI (Artificial Intelligence), 6. Performance, 7. Data, 8. Framework, 9. Reinforcement, 10. Applications) was reviewed by an expert from research area (B) “Data Analysis and Artificial Intelligence” and found to be potentially sensible to describe the field.

ChatGPT was also asked “How many publications published in 2023 can you detect in which 8 of the 10 terms of your previous answer are present?”; the answer was an elaboration of search strategies akin to what we describe in this paper, for example “You can search for combinations of the terms. For example, you can use boolean operators (AND, OR) to narrow down results.” (followed by an example).

At this point, we had used an alternative (LLM-based) method to derive terms; subsequently, these terms were used to construct a query.

Unfortunately, the search results (applied to area B and iterating through AND and OR combinations like before) were worse than those derived from our previous approaches, namely high FDR (>80%), low sensitivity (<5%) (cf. Table SI5b, columns B-D). Since not all PIs in area (B) “Data Analysis and Artificial Intelligence” are AI experts, we restricted the query to authors for which this is the case and repeated the best performing query. However, the sensitivity of the query was still very low (0.61%) as compared to the 50.91% of the query derived by the frequency-based analysis described above (Table SI5b, columne F-H).

These findings highlight the necessity of human oversight by experts and optimization of query structures, particularly when defining interdisciplinary research areas. While AI models might be useful in identifying relevant terms, a fully algorithmic query generation is currently unable to match the quality of a manually optimized search strategy, at least in the context of this publication.

## 4. Discussion

Three problems arise when trying to perform bibliometric analysis of trans- and interdisciplinary research: (1) selection of the “best” (balanced) query, (2) a delimitation problem and (3) the need for manual determination of TP and FP in order to evaluate the quality of the analysis.

(1) There is no clear statistical indicator which query is the “best”. Sensitivity and FDR of a query within a research area can be compared with each other, but it always remains the task of the analysing person to choose which combination of sensitivity and FDR and thus which query is feasible. With the metric used here, “(Sensitivity-FDR)+1”, we put equal weight on both characteristics. This evaluation method is nevertheless only possible for research areas which have a single and not dispersed affiliation. Furthermore, comparison of metrics between different research areas might be hampered by the different quality of queries. For example in area (C) a sensitivity of almost 84% with an FDR of 85% is the best result with respect to “|(Sensitivity – FDR)|”, while in area (B) only a maximum sensitivity of 56% could be achieved, with a simultaneously higher FDR of approx. 93%. A reason for this is the ease with which the various research areas can be captured. Research areas that are characterized by a high degree of homogenity, i.e., the authors use very similar vocabulary, are easier to define than research areas that draw on a very large and differentiated vocabulary. Thus, similar vocabluary leads to higher sensitivy and lower FDR values and vice versa (Figures S1-3).

Optimizing a query by optimizing the “(Sensitivity-FDR)+1” result still raises the question of the quality of the best query as the values for sensitivity and FDR are based on different populations. In addition, in some cases the values for FDR are in the order of the sensitivity and hence many FP will be detected. To get a better understanding of the quality of a query, we use an adaption of the ROC curve (receiver operating characteristics curve), a plot that illustrates the performance of a binary classifier model (Fawcett, 2006) and can be used for query balancing. For the application of a ROC-like curve in the context of the bibliometric analysis presented in this paper, it is important to keep in mind following aspects: For our analysis, it is not possible to determine TN and thus neither the specificity nor the false positive rate. Hence, we replaced specificity (x-axis) by FDR. In Figure SI1 the results in the context of such a modified ROC graph are depicted exemplatory for the queries with unigram terms for the research area (A), (B) and (C). As mentioned in the introduction, it is not possible to build a query which is perfect, i.e., a sensitivity of 100% and a FDR of 0%. Hence, the analysing person has to decide at which point the query is suitable for the question at hand. Another aspect for selecting the best query is the question of the interdisciplinarity of the retrieved publications. As the focus of this study is to specify and track interdisciplinary research areas, it is important to be able to retrieve interdisciplinary publications, which is not a trivial task. An analysis of a list of publications by looking at the title or abstract is usually sufficient if the person is knowledgeable about the research area in question, but in individual cases a deeper look might be required. If a query structure is based on terms (unigrams, bigrams or other types) which are linked with the OR operator only (see e.g. QUERY 5 with bigrams for the reseach area (D), the priority program), many disciplinary papers will be found. As already demonstrated with the bigrams for the priority program, it is in some cases unavoidable to use such query structures, as the absolute frequencies and distribution of the terms are too low among the publications, which makes it impossible to build more precice queries to retrieve interdisciplinary publications. Ideally, to get a higher fraction of interdisciplinary publications, the query structure should link some terms from different disciplines with the AND operator (see e.g. QUERY 3 and 4 with unigrams or QUERY 4.1 with bigrams in the above example). This can be applied only if the absolute frequency and distribution of the terms are high enough among the publications.

(2) Even if the selection of terms to be included in a query follows a clear concept, as described in the methodology section, the selection and combination of field tags and Boolean operators remains a “trial and error” procedure in which it is necessary to gradually move towards the optimal query. In particular, it is important to vary the selection of terms and to decide how many relevant terms should be included in the query. Too few terms lead to a limited coverage of the research area defined by the query, and too many terms may have the opposite effect and lead to the research area being poorly profiled or including overlapping research areas. The latter case in part can be solved by applying a NOT operator additionally and by selecting the terms (from FP publications) which thematically do not belong to the research area investigated. We suggest to start the selection of terms by using absolute frequencies (95% quartile), followed by using terms that occur with a relative frequency (in many abstracts) of 15% for high sensitivity of 30% for better specificity (cf. Methods and Figs. 1b/SI1). Such a rule of thumb is a good start, to be complemented where possible with expert-guided iteration of query formation and testing.

(3) If there exists no unique affiliation, as it is the case with e.g. priority programs, a calculation of the FDR is not possible without further definitions of FP and TP (see Methodology). To be able to do this, however, one must be both an expert in the research area and be able to invest time into the manual analysis. Hence, the person conducting the analysis needs to have a look at the literature output from a defined query and decide which publication is correct (TP) and which does not belong to the research area investigated (FP). This is additionally complicated by the fact that “correct” or “wrong” publications may depend on the context. In the strict sense, if defining for example a priority program, each publication that cannot be associated with one of its PIs, could be considered a FP. However, using such a strict definition, the research area as such cannot be defined for comparison with competitors on national/international level. Also, the simple association with a PI list will not work to define TPs, since PIs will publish in other research areas as well. Ultimately, the definition depends both on the research area investigated as well as which (bibliometric) questions the analysing person would like to answer. With the help of expert TP/FP declaration, the query can be refined iteratively to exclude those. This methodology can be applied fully only in cases where the number of publications retrieved from a query is not too high. For queries with high publication outputs, a random subset analysis might be feasible. As common in machine learning / classification approaches, several samples might be drawn and compared. A methodology of random subsampling and the subsequent iterative optimization of the query was applied by Elsevier for Sustainable Development Goals (SDG) mapping which are used for THE (Times Higher Education) Impact Rankings (Jayabalasingham et al., 2019). While such an approach is beyond the current scope of the study, it should be tested in future. Here, we show that random samples can be used to evaluate the performance of queries beyond the training data.

### Comparison to other methods

In addition to looking at the list of publications (i.e. thematical content of the publications) for judging whether they are indeed interdisciplinary, there are some other - both quantitative and qualitative - methods and approaches described in literature. First of all, it must be noted that in all these publications decribing these methods and approaches, it is highly debated what the definition of interdisciplinarity is which in turn affects the methodology to be applied and the interpretation of its results.

According to (Glänzel & Debackere, 2022) there are two different concepts for classifying interdisciplinarity, i.e. the *cognitive approach* and the *organisational approach*. The cognitive approach considers aspects such as information flows (e.g. citations) and the organisational approach investigates the collaboration (e.g. co-authorship based on affiliation and subject classification of the researcher). In the framework of these two concepts, there are two subsequent main concepts which are state of the art in bibliometrics, the concepts of *diversity* and *coherence*. The concept diversity consists of three indicators: *variety* (how many disciplines are involved), *balance* (how much do each of these disciplines contribute) and *disparity* (how different from each other are the involved disciplines). The concept coherence considers knowledge integration, i.e. the analysis to what extent topics, tools, data and other aspects of scientific content from different disciplines contribute and result in the research. Thus, the concept diversity deals with the disciplinary constituents of the research and coherence with the cooperation and networking leading to new results, ideas, concepts, tools, etc. These concepts inevitably lead to different definitions, qualitative and quantitative methods and interpretation of interdisciplinarity. It should also be noted that approaches that try to determine the interdisciplinary of the research conducted by a given researcher might be misleading, since researchers will often publish in disciplinary as well as (several) interdisciplinary research areas.

How would the methods outlined above perform on the data used here? In case of the priority program (D) the training set of publications allows to identify the following areas mentioned before: Plant Sciences (26), Cell Biology (5), Biochemistry & Molecular Biology (4), Biology (3), Reproductive Biology (2), Developmental Biology (1), Ecology (1), Evolutionary Biology (1), Genetics & Heredity (1) and Multidisciplinary Sciences (1). With these statistics we can derive a qualitative judgement based on the diversity concept about how many disciplines are involved (variety), how much do each of these disciplines contribute (balance) and how different from each other the involved disciplines are (disparity). Depending on a strict or loose definition of interdisciplinarity (here with respect to e.g. diversity and disparity of the contributing subjects) these sets of publications can be regarded interdisciplinary. This evaluation of the training set could serve as a reference for analysing a list of publication by a query based on the same training set. However, this would only give the information of whether a publication can be considered interdisciplinary, and cannot be used to assign it to an interdisciplinary research area. Moreover, as pointed out above, the use of disciplinary terms can be highly misleading.

### Caveats

The method presented here relies on a set of training data that stem from the interdisciplinary research area in question. Publications in the training data that are selected for the wrong reasons (for example journal impact rather than topic) will lead to a less efficient selection of appropriate terms.

The iterative improvement of queries can to some extent be performed by a person knowledgeable in the methodology (bibliometric term analyses), but will benefit from expert knowledge on the interdisciplinary research area. Similarly, analysis of results beyond the training data, e.g. the random sample analyses used here, will require expert knowledge to determine TP.

## 5. Conclusion

The term-based bibliometric analysis method introduced here requires a TP “training set” to start with. Given that, and a willingness to develop and test several queries, this method can be used to define interdisciplinary research areas with moderate accuracy (good sensitivity at high FDR). A rule of thumb for term selection based on absolute (95% quartile) and relative (15%) term frequencies may be used if expert knowledge on the research field is not available.

Unigrams have a higher sensitivity due to their more general nature and higher absolute and relative frequencies in publications, but lead to a higher FP rate. The use of bigrams avoids some of the FP problem of unigrams, but care needs to be taken that they do not drastically reduce sensitivity if they are a required argument (AND) and appear with low frequency in published work of the area. The exclusion of terms via a NOT statement is a good option to reduce FP without losing too many TP.

A term-based bibliometric analysis may be used to define interdisciplinary research areas and to subsequently track their development, for example within an institution (how many publications, by whom and considering a time course), or between institutions (national or international ranking, detection of institutions that publish in this field).

## Acknowledgements

We would like to thank the members of the research areas studied here, and two anonymous reviewers of a previous version of the manuscript for providing valuable input. This project was carried out using institutional funding of the University of Freiburg, and in the framework of MAdLand (https://madland.science, DFG priority programme 2237), SAR is grateful for funding by the DFG (RE-1697/19-1).

## Supplementary Information (SI)

Table SI1: Raw data of the four example research areas on which the study was conducted.

Table SI2: Table of research area D/MAdLand publications and which of them are detected by which query.

Table SI3: Research area (D) QUERY 2 (“better sensitivity”) vs. MAdLand QUERY 4 (“better specificity”) at the level of WoS Citation Topics Meso.

Table SI4: List of unigram/bigram NOT operator terms to reduce medical and other off-topic literature in research area (D) MAdLand.

Table SI5: Large language model approach to define AI publications and terms (A) and related queries (B).

Figure SI1: Absolute and relative frequencies of terms derived from the training data set word count analysis for the research areas (A)-(D).

Figure SI2: ROC-like diagram for the queries with unigrams for the research areas (A), (B) and (C) for QUERY 1-4.

Figure SI3: Sensitivity and False Discovery Rate (FDR) and (Sens-FDR)+1 of the research area (B) “Data Analysis and Artificial Intelligence”.

Figure SI4: Sensitivity and False Discovery Rate (FDR) and (Sens-FDR)+1 of the research area (A) “Epigenetics, Immunology and Cancer Research”.

