## Supplementary material for "How bibliometric analysis can help to specify and track interdisciplinary research areas: A term-based analysis approach": Figures SI1-SI4, Tables SI3+SI4

### Slide 1
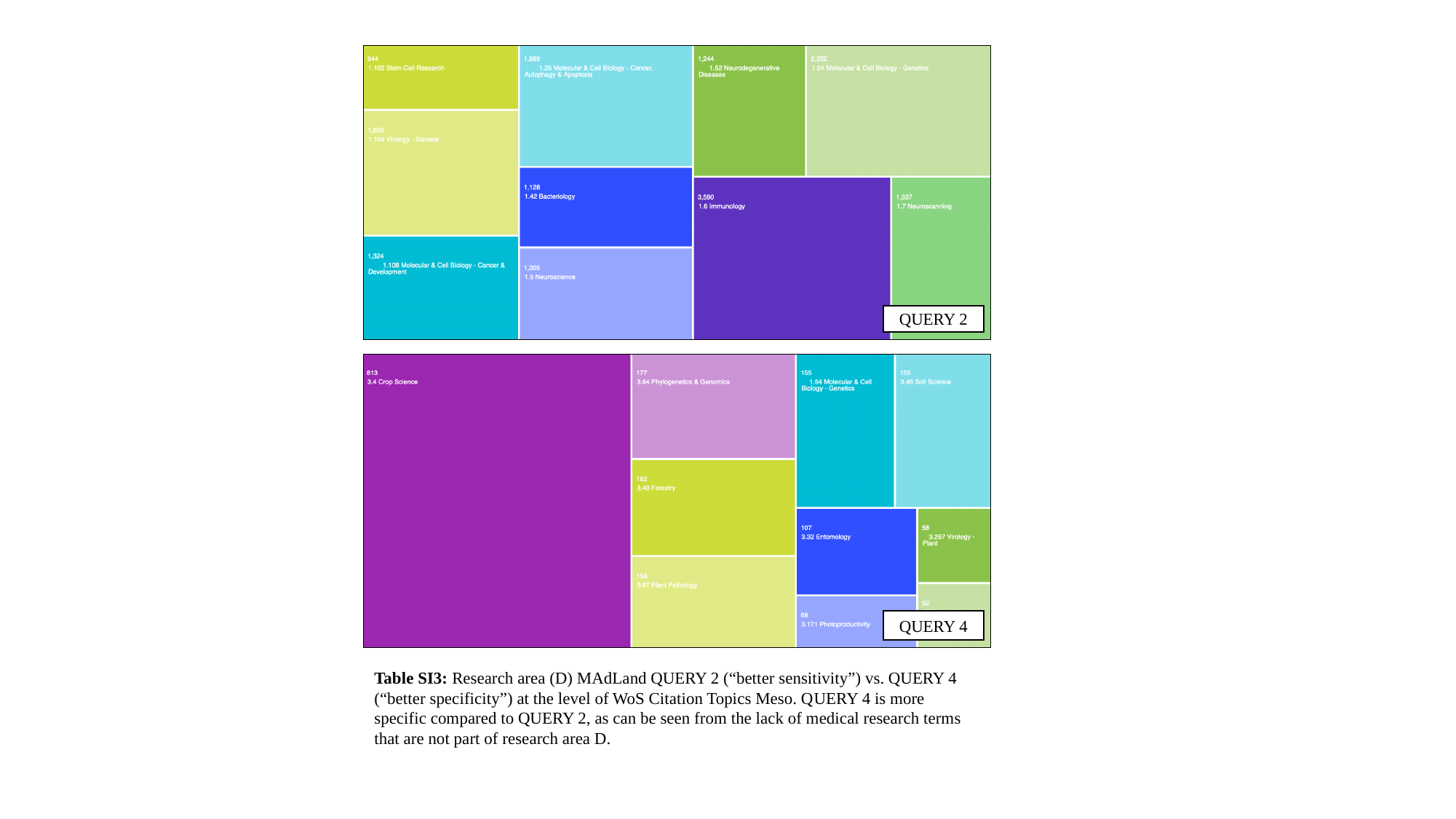

QUERY 2
QUERY 4
Table SI3: Research area (D) MAdLand QUERY 2 (“better sensitivity”) vs. QUERY 4 (“better specificity”) at the level of WoS Citation Topics Meso. QUERY 4 is more specific compared to QUERY 2, as can be seen from the lack of medical research terms that are not part of research area D.

### Slide 2
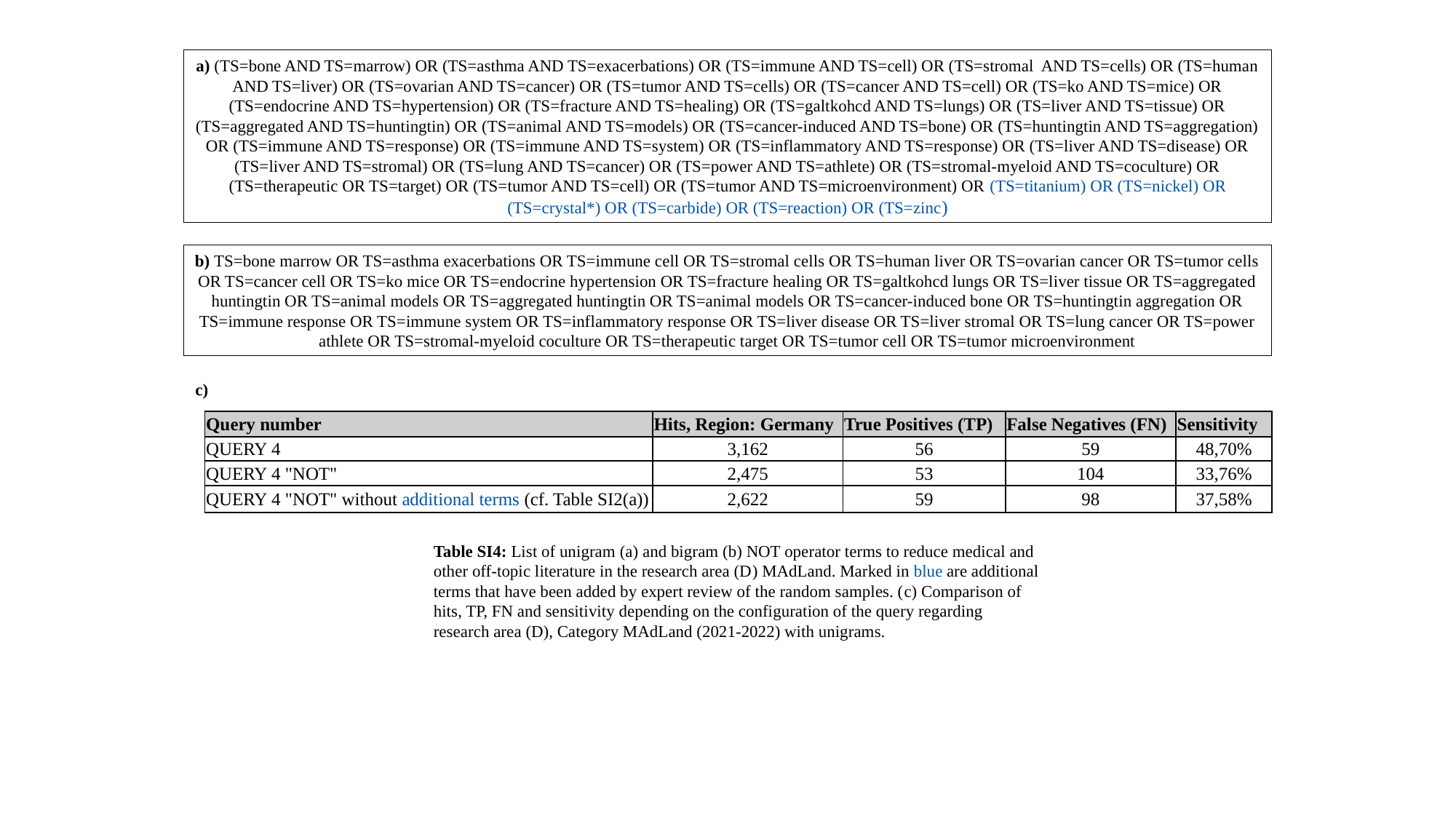

a) (TS=bone AND TS=marrow) OR (TS=asthma AND TS=exacerbations) OR (TS=immune AND TS=cell) OR (TS=stromal AND TS=cells) OR (TS=human AND TS=liver) OR (TS=ovarian AND TS=cancer) OR (TS=tumor AND TS=cells) OR (TS=cancer AND TS=cell) OR (TS=ko AND TS=mice) OR (TS=endocrine AND TS=hypertension) OR (TS=fracture AND TS=healing) OR (TS=galtkohcd AND TS=lungs) OR (TS=liver AND TS=tissue) OR (TS=aggregated AND TS=huntingtin) OR (TS=animal AND TS=models) OR (TS=cancer-induced AND TS=bone) OR (TS=huntingtin AND TS=aggregation) OR (TS=immune AND TS=response) OR (TS=immune AND TS=system) OR (TS=inflammatory AND TS=response) OR (TS=liver AND TS=disease) OR (TS=liver AND TS=stromal) OR (TS=lung AND TS=cancer) OR (TS=power AND TS=athlete) OR (TS=stromal-myeloid AND TS=coculture) OR (TS=therapeutic OR TS=target) OR (TS=tumor AND TS=cell) OR (TS=tumor AND TS=microenvironment) OR (TS=titanium) OR (TS=nickel) OR (TS=crystal*) OR (TS=carbide) OR (TS=reaction) OR (TS=zinc)
b) TS=bone marrow OR TS=asthma exacerbations OR TS=immune cell OR TS=stromal cells OR TS=human liver OR TS=ovarian cancer OR TS=tumor cells OR TS=cancer cell OR TS=ko mice OR TS=endocrine hypertension OR TS=fracture healing OR TS=galtkohcd lungs OR TS=liver tissue OR TS=aggregated huntingtin OR TS=animal models OR TS=aggregated huntingtin OR TS=animal models OR TS=cancer-induced bone OR TS=huntingtin aggregation OR TS=immune response OR TS=immune system OR TS=inflammatory response OR TS=liver disease OR TS=liver stromal OR TS=lung cancer OR TS=power athlete OR TS=stromal-myeloid coculture OR TS=therapeutic target OR TS=tumor cell OR TS=tumor microenvironment
Table SI4: List of unigram (a) and bigram (b) NOT operator terms to reduce medical and other off-topic literature in the research area (D) MAdLand. Marked in blue are additional terms that have been added by expert review of the random samples. (c) Comparison of hits, TP, FN and sensitivity depending on the configuration of the query regarding research area (D), Category MAdLand (2021-2022) with unigrams.
c)
| Query number | Hits, Region: Germany | True Positives (TP) | False Negatives (FN) | Sensitivity |
| --- | --- | --- | --- | --- |
| QUERY 4 | 3,162 | 56 | 59 | 48,70% |
| QUERY 4 "NOT" | 2,475 | 53 | 104 | 33,76% |
| QUERY 4 "NOT" without additional terms (cf. Table SI2(a)) | 2,622 | 59 | 98 | 37,58% |

### Slide 3
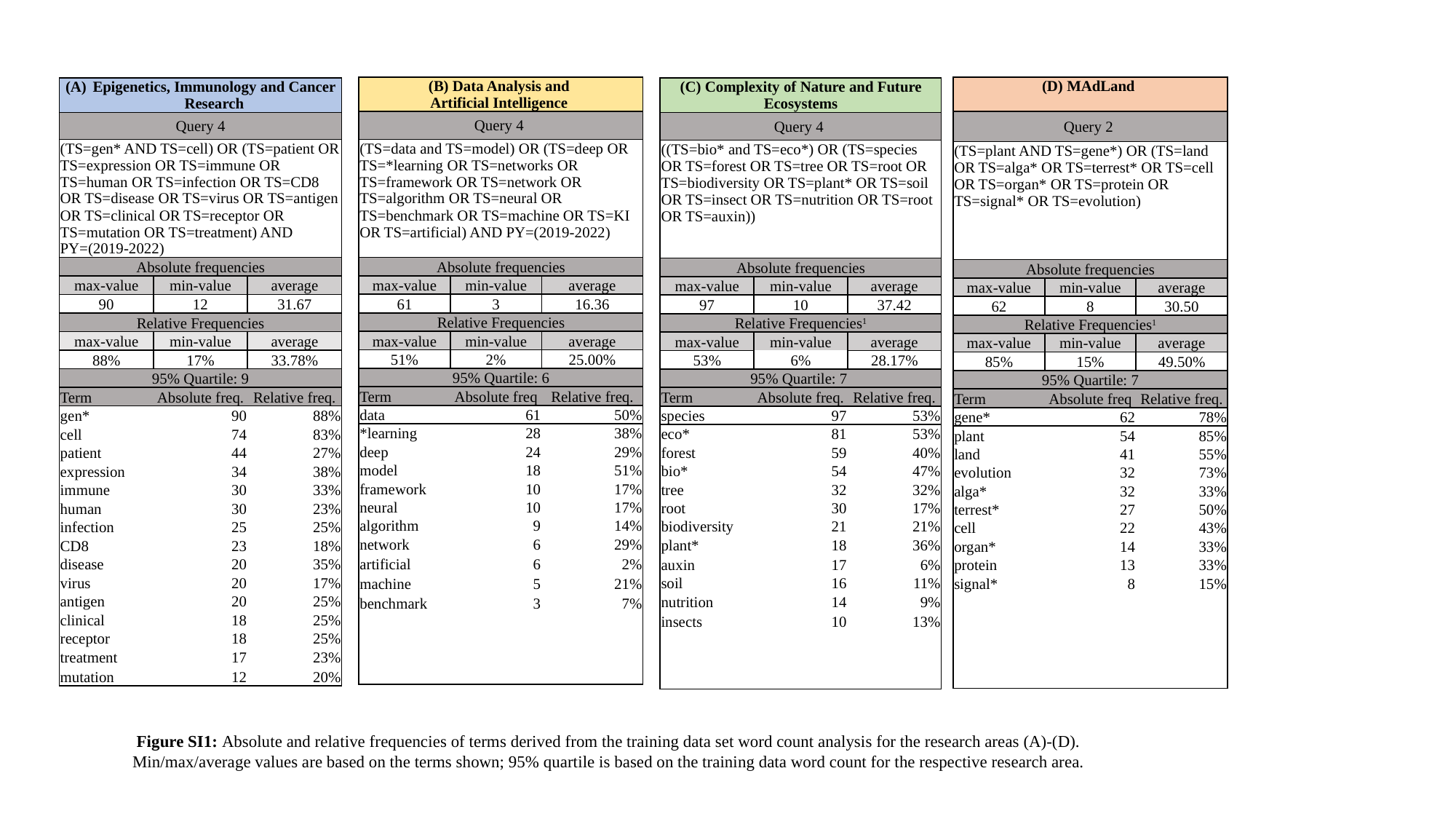

| (B) Data Analysis and Artificial Intelligence | | |
| --- | --- | --- |
| Query 4 | | |
| (TS=data and TS=model) OR (TS=deep OR TS=\*learning OR TS=networks OR TS=framework OR TS=network OR TS=algorithm OR TS=neural OR TS=benchmark OR TS=machine OR TS=KI OR TS=artificial) AND PY=(2019-2022) | | |
| Absolute frequencies | | |
| max-value | min-value | average |
| 61 | 3 | 16.36 |
| Relative Frequencies | | |
| max-value | min-value | average |
| 51% | 2% | 25.00% |
| 95% Quartile: 6 | | |
| Term | Absolute freq | Relative freq. |
| data | 61 | 50% |
| \*learning | 28 | 38% |
| deep | 24 | 29% |
| model | 18 | 51% |
| framework | 10 | 17% |
| neural | 10 | 17% |
| algorithm | 9 | 14% |
| network | 6 | 29% |
| artificial | 6 | 2% |
| machine | 5 | 21% |
| benchmark | 3 | 7% |
| (D) MAdLand | | |
| --- | --- | --- |
| Query 2 | | |
| (TS=plant AND TS=gene\*) OR (TS=land OR TS=alga\* OR TS=terrest\* OR TS=cell OR TS=organ\* OR TS=protein OR TS=signal\* OR TS=evolution) | | |
| Absolute frequencies | | |
| max-value | min-value | average |
| 62 | 8 | 30.50 |
| Relative Frequencies^1^ | | |
| max-value | min-value | average |
| 85% | 15% | 49.50% |
| 95% Quartile: 7 | | |
| Term | Absolute freq | Relative freq. |
| gene\* | 62 | 78% |
| plant | 54 | 85% |
| land | 41 | 55% |
| evolution | 32 | 73% |
| alga\* | 32 | 33% |
| terrest\* | 27 | 50% |
| cell | 22 | 43% |
| organ\* | 14 | 33% |
| protein | 13 | 33% |
| signal\* | 8 | 15% |
| Epigenetics, Immunology and Cancer Research | | |
| --- | --- | --- |
| Query 4 | | |
| (TS=gen\* AND TS=cell) OR (TS=patient OR TS=expression OR TS=immune OR TS=human OR TS=infection OR TS=CD8 OR TS=disease OR TS=virus OR TS=antigen OR TS=clinical OR TS=receptor OR TS=mutation OR TS=treatment) AND PY=(2019-2022) | | |
| Absolute frequencies | | |
| max-value | min-value | average |
| 90 | 12 | 31.67 |
| Relative Frequencies | | |
| max-value | min-value | average |
| 88% | 17% | 33.78% |
| 95% Quartile: 9 | | |
| Term | Absolute freq. | Relative freq. |
| gen\* | 90 | 88% |
| cell | 74 | 83% |
| patient | 44 | 27% |
| expression | 34 | 38% |
| immune | 30 | 33% |
| human | 30 | 23% |
| infection | 25 | 25% |
| CD8 | 23 | 18% |
| disease | 20 | 35% |
| virus | 20 | 17% |
| antigen | 20 | 25% |
| clinical | 18 | 25% |
| receptor | 18 | 25% |
| treatment | 17 | 23% |
| mutation | 12 | 20% |
| (C) Complexity of Nature and Future Ecosystems | | |
| --- | --- | --- |
| Query 4 | | |
| ((TS=bio\* and TS=eco\*) OR (TS=species OR TS=forest OR TS=tree OR TS=root OR TS=biodiversity OR TS=plant\* OR TS=soil OR TS=insect OR TS=nutrition OR TS=root OR TS=auxin)) | | |
| Absolute frequencies | | |
| max-value | min-value | average |
| 97 | 10 | 37.42 |
| Relative Frequencies^1^ | | |
| max-value | min-value | average |
| 53% | 6% | 28.17% |
| 95% Quartile: 7 | | |
| Term | Absolute freq. | Relative freq. |
| species | 97 | 53% |
| eco\* | 81 | 53% |
| forest | 59 | 40% |
| bio\* | 54 | 47% |
| tree | 32 | 32% |
| root | 30 | 17% |
| biodiversity | 21 | 21% |
| plant\* | 18 | 36% |
| auxin | 17 | 6% |
| soil | 16 | 11% |
| nutrition | 14 | 9% |
| insects | 10 | 13% |
Figure SI1: Absolute and relative frequencies of terms derived from the training data set word count analysis for the research areas (A)-(D).
Min/max/average values are based on the terms shown; 95% quartile is based on the training data word count for the respective research area.

### Slide 4
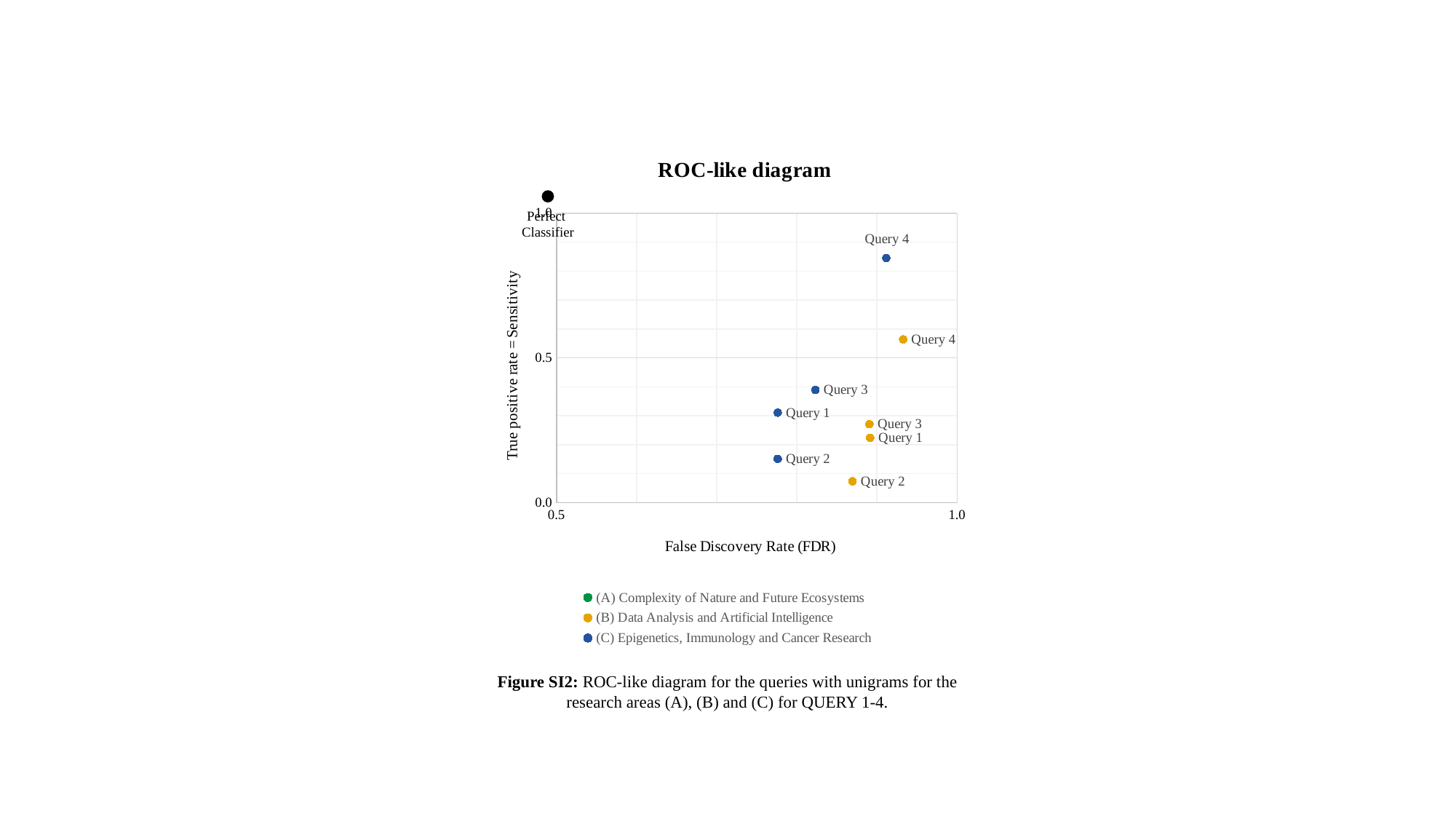

#### Chart: ROC-like diagram
| Category |
|---|
Perfect
Classifier
Figure SI2: ROC-like diagram for the queries with unigrams for the research areas (A), (B) and (C) for QUERY 1-4.

### Slide 5
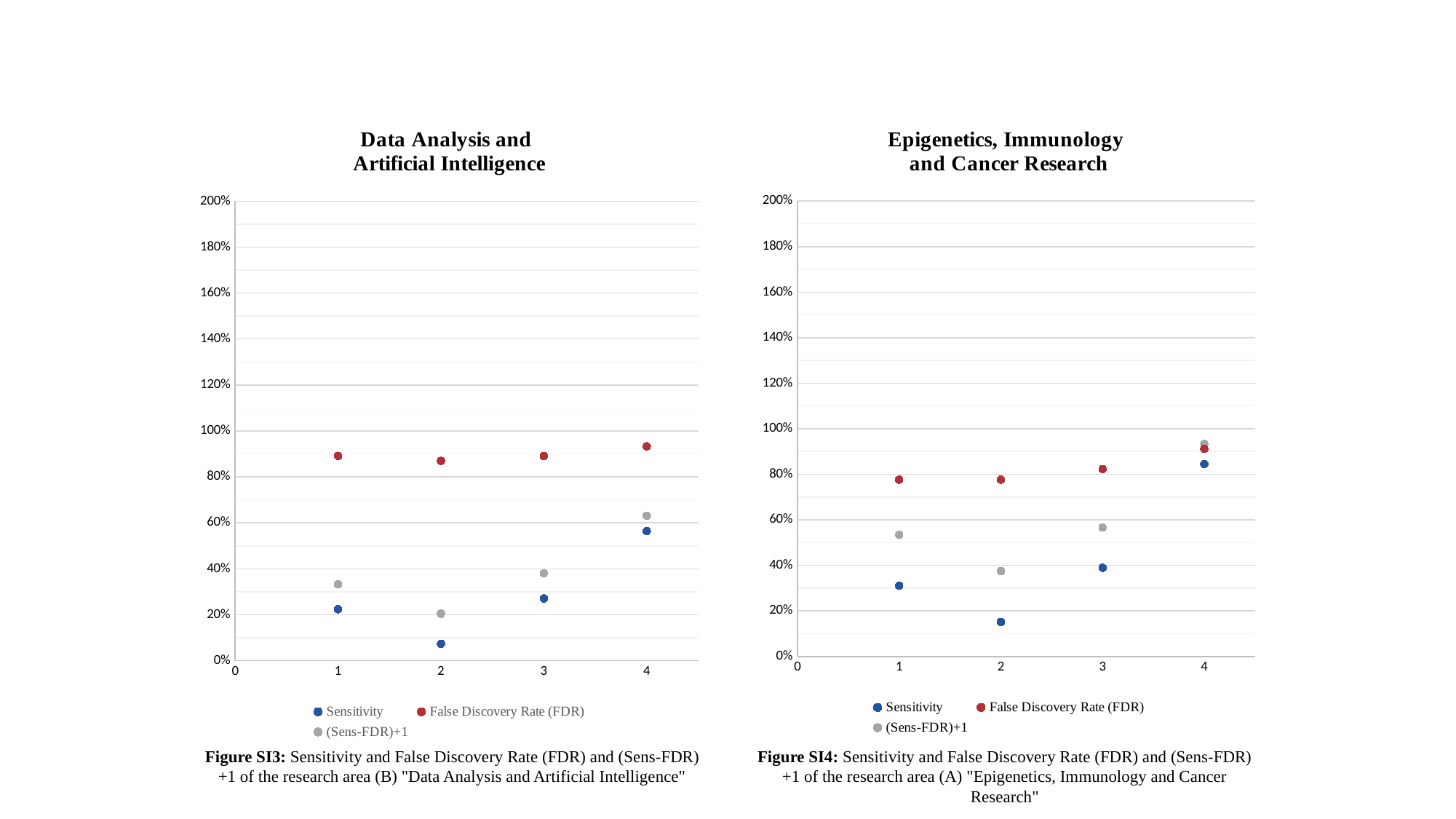

#### Chart: Data Analysis and
Artificial Intelligence
| Category | Sensitivity | False Discovery Rate (FDR) | (Sens-FDR)+1 |
|---|---|---|---|
#### Chart: Epigenetics, Immunology
and Cancer Research
| Category | Sensitivity | False Discovery Rate (FDR) | (Sens-FDR)+1 |
|---|---|---|---|Figure SI3: Sensitivity and False Discovery Rate (FDR) and (Sens-FDR)+1 of the research area (B) "Data Analysis and Artificial Intelligence"
Figure SI4: Sensitivity and False Discovery Rate (FDR) and (Sens-FDR)+1 of the research area (A) "Epigenetics, Immunology and Cancer Research"
